# Non-invasive visualization of amyloid-beta deposits in Alzheimer amyloidosis mice using magnetic resonance imaging and fluorescence molecular tomography

**DOI:** 10.1101/2021.11.20.466221

**Authors:** Wuwei Ren, Ruiqing Ni, Markus Vaas, Jan Klohs, Jorge Ripoll, Martin Wolf, Markus Rudin

## Abstract

Abnormal cerebral accumulation of amyloid-beta peptide (Aβ) is a major hallmark of Alzheimer’s disease. Non-invasive monitoring of Aβ deposits enables assessing the disease burden in patients and animal models mimicking aspects of the human disease as well as evaluating the efficacy of Aβ-modulating therapies. Previous *in vivo* assessments of plaque load in mouse models of cerebral amyloidosis have been predominantly based on two-dimensional diffuse fluorescence reflectance imaging (2D-FRI) and two-photon microscopy (2PM) using Aβ-specific imaging agents. However, 2D-FRI lacks depth resolution, whereas 2PM is restricted by the limited field of view preventing coverage of large brain regions. Here, we utilized a magnetic resonance imaging (MRI) and fluorescence molecular tomography (FMT) pipeline with the curcumin derivative fluorescent probe CRANAD-2 to achieve full 3D brain coverage for detecting Aβ accumulation in the arcAβ mouse model of cerebral amyloidosis. A homebuilt FMT system was used for data acquisition in combination with a customized software platform enabling the integration of anatomical information derived from MRI as prior information for FMT image reconstruction. The results obtained from the FMT-MRI study were compared to data obtained from conventional 2D-FRI recorded under similar physiological conditions. The two methods yielded comparable time courses of the fluorescence intensity following intravenous injection of CRANAD-2 in a region of interest comprising the mouse brain. The depth resolution inherent to FMT allowed separation of signal contributions from the scalp and different brain regions, indicating preferential accumulation of the fluorescent tracer in the cerebral cortex, a region characterized by significant plaque deposition in arc Aβ mice. In conclusion, we have demonstrated the feasibility of visualizing Aβ deposition in 3D using a multimodal FMT-MRI method. This hybrid imaging method provides complementary anatomical, physiological and molecular information, thereby enabling the detailed characterization of the disease status in mouse models of cerebral amyloidosis, which is also important for monitoring the efficacy of putative treatments targeting Aβ.

## 1 Introduction

The abnormal accumulation and spread of amyloid-beta (Aβ) deposits plays a central role in the pathogenesis of Alzheimer’s disease (AD), leading to neurotoxicity and inflammation and synaptic dysfunction (De Strooper and Karran, 2016; Walsh et al., 2002; Zott et al., 2019). Aβ assembles into oligomer, fibrillar Aβ and Aβ plaques (Cohen et al., 2013) in a spontaneous manner. Positron emission tomography (PET) imaging with amyloid tracers such as [^11^C]PIB (Klunk et al., 2005), [^18^F]florbetaben (Barthel et al., 2011) and [^18^F]florbetapir (Fleisher et al., 2011) has shown higher cortical fibrillar Aβ loads in patients with AD continuum compared to cognitive normal controls (Ni et al., 2017; Villemagne et al., 2018). Amyloid PET imaging has been promoted as a pathological imaging biomarker for the early and differential diagnosis of AD and has demonstrated clinical utility (Jack et al., 2018).

In preclinical research, *in vivo* Aβ detection and longitudinal monitoring in animal models of AD amyloidosis provided a handle for studying the effect of targeted molecular interventions on this molecular phenotype, thereby providing indirect insight into the mechanism underlying Aβ deposition and potential biomarkers for assessing the effects of Aβ-modulating therapeutics (Ni, 2021b). Detection has been made possible by using imaging tools covering different ranges of spatial resolution and fields-of-view (FOV) (Razansky et al., 2021). At the macroscopic level, three-dimensional (3D) microPET with [^11^C]PIB (Snellman et al., 2017), [^18^F]florbetapir (Poisnel et al., 2012), [^18^F]florbetaben (Yousefi et al., 2015), [^18^F]flutemetamol, [^11^C]AZD2184 (Rodriguez-Vieitez et al., 2015), and [^124^I]labelled antibody (Sehlin et al., 2016) has enabled sensitive detection of Aβ deposits for longitudinal imaging. Imaging of Aβ deposits at a mesoscopic resolution has also been demonstrated using ^19^F- and ^1^H-magnetic resonance imaging (MRI) with or without contrast agents (Higuchi et al., 2005; Ni, 2021a) or u using combined MRI - near-infrared fluorescence (NIRF) imaging in combination with dual modal NIRF-MRI probes (Li et al., 2018). The development of Aβ-specific fluorescent probes such as AOI987 (Hintersteiner et al., 2005), CRANAD-2/-3 (Ran et al., 2009; Zhang et al., 2015), and luminescent conjugated oligothiophenes (Calvo-Rodriguez et al., 2019; Ni et al., 2020a; Shirani et al., 2015) using methoxy-X04, BTA-1 and PIB (Bacskai et al., 2003; Hefendehl et al., 2011; Klunk et al., 2002; Meyer-Luehmann et al., 2008) enabled the application of various optical imaging methods in preclinical studies. Optical detection of Aβ deposits by using *ex vivo* optical projection tomography (Nguyen et al., 2019), light-sheet microscopy (Ni et al., 2020b) and various *in vivo* fluorescence microscopy methods, such as multiphoton microscopy, enabled monitoring of Aβ at *μ*m spatial resolution (Pansieri et al., 2019). However, two-photon imaging techniques hampered by a limited field-of-view (FOV) and due to light scattering effects commonly involve cranial opening to expose the brain surface, which may affect brain physiology.

Compared with the aforementioned optical imaging methods, fluorescence molecular tomography (FMT) is a truly non-invasive imaging tool that covers a large FOV comparable to traditional fluorescence reflectance imaging (FRI) and resolves the fluorophore distribution in tissue in 3D (Ntziachristos et al., 2002). More specifically, FMT belongs to a subclass of diffuse optical imaging modalities that adopt near-infrared light for exciting the fluorescence probe and utilize a model-based reconstruction algorithm to recover the 3D distribution of the fluorophore (Arridge and Schotland, 2009). In the last decade, FMT has been applied widely in preclinical settings focusing predominantly on early tumor detection and monitoring (Darne et al., 2014; Liu et al., 2015). In the study of AD, a multimodal imaging method that combines FMT and X-ray computer tomography (CT) has been applied to map the distribution of Aβ plaques employing the oxazine derivative AO1987 probe in a static manner (Hyde et al., 2009). The multimodal strategy accounts for the loss of structural information inherent to most molecular imaging techniques, including FMT and PET. Registration of molecular imaging data to anatomical reference image recorders under identical conditions is of value for predefined volume-of-interest (VOI)-based quantitative data analysis. Compared with FMT-CT, the combination of FMT and MRI has several advantages: MRI provides better image contrast on soft tissue, especially in the field of brain imaging. In addition, both clinical and preclinical high-field MRI scanners can offer versatile physiological (e.g., regional cerebral blood flow, blood-brain barrier integrity, cerebral oxygenation) functional (e.g., functional neuronal networks) and structural information (e.g., white matter integrity, brain atrophy). Thus, the FMT-MRI combination allows the spatial linkage between the functional/structural (MRI) and molecular information (FMT). FMT-MRI has been applied to study aspects of tumor vascularization in a mouse model (Ren et al., 2016b). To the best of our knowledge, the hybrid approach has not yet been applied for studying cerebral pathologies such as cerebral amyloidosis.

In the present study, we demonstrated whole-brain mapping of Aβ deposits at mm resolution in arcAβ mouse models of cerebral amyloidosis, as observed in AD (Merlini et al., 2011), using the Aβ fibril binding curcumin derivative CRANAD-2 as a fluorescent reporter. The *in vivo* imaging results were validated by immunofluorescence staining using CRANAD-2 with the anti-Aβ antibody 6E10 on mouse brain sections.

## 2 Material and methods

### 2.1 Animal models

Five transgenic arcAβ mice overexpressing the human APP695 transgene containing the Swedish (K670N/M671L) and Arctic (E693G) mutations under the control of the prion protein promoter and five age-matched nontransgenic littermates of both sexes were used in this study (18-20 months of age) (Knobloch et al., 2007; Merlini et al., 2011; Ni et al., 2019; Ni et al., 2018). Animals were housed in ventilated cages inside a temperature-controlled room and under a 12-hour dark/light cycle. Pelleted food (3437PXL15, CARGILL) and water were provided ad libitum. All experiments were performed in accordance with the Swiss Federal Act on Animal Protection and were approved by the Cantonal Veterinary Office Zurich (permit number: ZH018/14). All procedures fulfilled the ARRIVE guidelines on reporting animal experiments.

### 2.2 Preparation of animals and fluorescence probe

Briefly, mice were first anesthetized with an initial dose of 4 % isoflurane (Abbott, Cham, Switzerland) in an oxygen/air mixture (200/800 mL/min), and anesthesia was maintained at 1.5 % isoflurane in oxygen/air (100/400 mL/min). The fur over the head was then removed. The mice were placed in the prone position on a heating pad that maintained a constant body temperature (36.5 degrees). Regarding the fluorescent reporter, we used the Aβ-specific curcumin derivative CRANAD-2 (Sigma, Switzerland) (Ran et al., 2009) dissolved in a buffer of 20 % dimethyl sulfoxide/polyethylene glycol (DMSO/PEG, Sigma, Switzerland) for *in vivo* imaging of amyloid aggregates in arcAβ mice. The spectrum of CRANAD-2 shows an absorption maximum at 640 nm and an emission maximum at 715 nm upon interacting with Aβ aggregates, undergoing a 90 nm blueshift from 805 nm in the absence of Aβ. Moreover, the target interaction has been reported to result in a large increase in the quantum yield.

### 2.3 Magnetic resonance imaging

MRI scans were performed on a 7 T small animal MRI BioSpec scanner (Bruker Biospin GmbH, Ettlingen, Germany) with a magnet bore diameter of 16 cm and equipped with an actively shielded gradient capable of switching 760 mT/ m with an 80-μs rise time and operated by the ParaVision 6.0 software platform (Bruker Biospin GmbH, Ettlingen, Germany). A circular polarized volume resonator was used for signal transmission, and an actively decoupled mouse brain quadrature surface coil with integrated combiner and preamplifier was used for signal reception. *In vivo* T_2_*-weighted MR images of mouse brain/head were recorded using a gradient echo sequence using the following parameters: FOV 20 × 20 mm^2^, number of slices 35, dimension of reconstructed matrix 256 × 256 × 35, slice thickness 0.5 mm, repetition time 4200 ms, echo time 33 ms, and flip angle α 90 degree within a scan time 12 min 36 s.

### 2.4 Fluorescence molecular tomography

A homebuilt non-contact FMT system was used for data acquisition (Ren et al., 2020b). The system is equipped with a 16-bit charge-coupled device (CCD) camera (ANDOR, Belfast, Northern Ireland), a galvanometric driven mirror system (ScanLab, Puchheim, Germany), a solid-state laser generator with 670 nm wavelength for illumination (B&W Tek, Newark, USA), and custom-made animal support. The camera was cooled to -79 degrees for noise reduction. Before brain imaging, the fur overlying the head was first trimmed with an electrical shaver and then removed using a depilation cream (Nair, NJ, USA). The tail vein was then cannulated with a 30-gauge needle for subsequent intravenous injection with 2 mg/kg CRANAD-2 solution in 20% DMSO/PEG buffer. The illumination pattern was designed as a 7 × 7 grid covering the mouse head. Bandpass filters with central frequencies of 660 ± 13 nm and 720 ± 13 nm for excitation and emission procedures, respectively, were used. Measurements were carried out prior to and 5 min, 10 min, 20 min, and 30 min after injection of CRANAD-2. To ensure continuous recording of changes in the fluorescence signal, only one excitation image and one white-light image were taken prior to label injection, while emission images were taken at the four time points as indicated before.

### 2.5 FMT-MRI data processing

We used our customized reconstruction platform to perform FMT-MRI image reconstruction (Ren et al., 2020a). The schematics of FMT-MRI data processing are shown schematically in **Fig. 1**. First. The T_2_*-weighted MR images were used as the input, and the white-light image was registered to the topological MRI map, which represents the mouse surface, using a landmark-based registration method. With common coordinates established, a self-adaptive mesh was generated based on the MRI anatomy (Ren et al., 2016a). The values of absorption and scattering coefficients were also generated according to the intensity-based segmentation of the MR images using published values of optical properties for different biological tissue types (Jacques, 2013). For simplicity, only four tissue categories were considered (**Table. 1**). The refractive index was set uniformly to n = 1.4. The virtual detector plane consisting of individual elements with an area of 1×1 mm^2^ was assigned to the surface of the mesh. The diameter of each laser point was set to 1 mm. To merge the FMT-MRI images and analyse the results, a linear interpolation scheme was used to transform from the mesh coordinates used for reconstructing the dye distribution to Cartesian grid coordinates of the anatomical reference image. A Tikhonov regularization term was introduced to the reconstruction, and the inverse problem was solved using the conjugation gradient (CG) method (Ren et al., 2020a; Ren et al., 2020b). Quantitative analysis was carried out following FMT reconstruction. The registered MR images provided an anatomical reference for defining 3D VOIs using the Allen Brain Atlas as a reference (Jones et al., 2009). We analysed the temporal variation in the cortical and subcortical regions. The average signal intensities within these two VOIs were calculated preinjection and 5 min, 10 min, 20 min, and 30 min after injection. The uptake curves of CRANAD-2 in cortical and subcortical regions are depicted.

**Figure 1:**
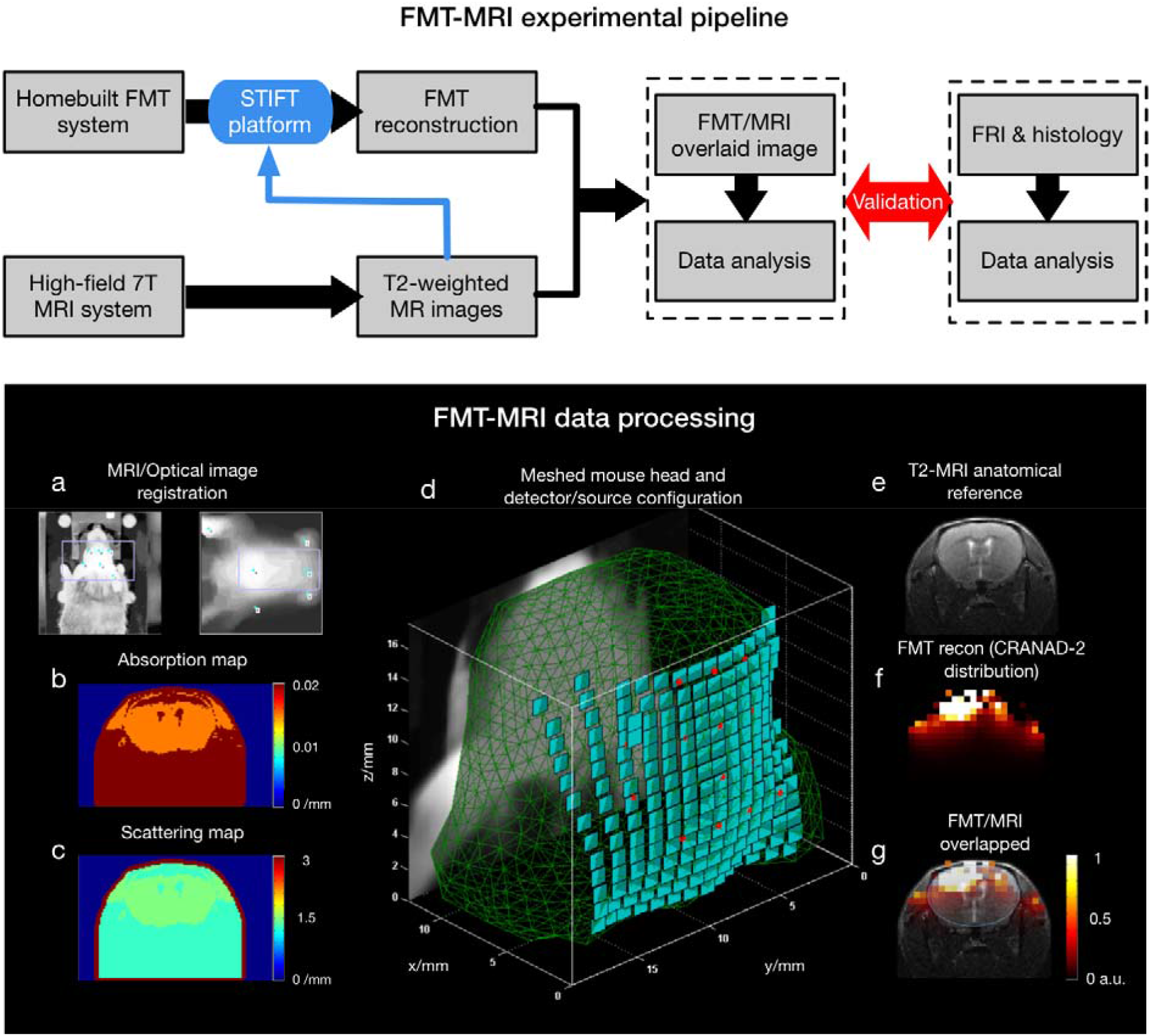
The multimodal FMT-MRI experimental pipeline and data processing framework are illustrated in the upper and lower panels, respectively. Regarding the framework of FMT-MRI data processing, the topological surface of anatomical MRI (a, right) was first registered with the optical image (a, left) with fiducial markers indicated on both images (blue dot). Once the two types of images were registered, both an absorption map (b) and a scattering map (c) were generated according to the intensity-based segmentation. (d) MRI-derived structural information was used for reconstructing the object in 3D, and adaptive meshing was performed with detector elements (green patches) and laser illumination points (red spots) assigned to the surface of the generated mesh. (e) The T_2_-weighted MR image (e) provides an important anatomical reference that can significantly improve the localization of the FMT signal (f). (g) shows an overlaid FMT-MRI image containing both molecular and structural information within the same mouse.

### 2.6 Comparison of FMT and fluorescence reflectance imaging

FRI, also referring to near-infrared fluorescence imaging, was performed for comparison with the results obtained from FMT. We used the Maestro 500 multispectral imaging system (CRi, MA, USA) (Klohs et al., 2009). Briefly, the imager was equipped with a bandpass filter (615 to 665 nm) for excitation. The fluorescence signal was detected by a CCD camera mounted on the top of the imaging chamber. Images were acquired by increasing the emission filter central wavelengths by 10-nm increments. Image cubes recorded were spectrally unmixed using Maestro software (CRi, MA, USA).

### 2.7 Immunofluorescence staining and confocal microscopy

For staining and immunofluorescence investigations, mice were perfused with 1 × PBS (pH 7.4) under ketamine/xylazine/acepromazine maleate anesthesia (75/10/2 mg/kg body weight, bolus injection) and decapitated. Brains were removed from the skull afterwards and fixed in 4% paraformaldehyde in 1 × PBS (pH 7.4). Brain hemispheres were embedded in paraffin following routine procedures and cut into 5 μm horizontal sections. Immunofluorescence staining using donkey-anti-rat Alexa488 (Jackson, AB-2340686, 1:400), anti-Aβ_1–16_ antibody 6E10 (Signet Lab, SIG-39320, 1:5000), thioflavin S, and CRANAD-2 (Kayed et al., 2007) was performed following a protocol described earlier with nuclei counterstained by 4’ 6-diamidino-2-phenylindole (DAPI, Sigma, D9542 10MG, 1:1000). To further assess the colocalization of different channels, confocal images of arcAβ mice and non-transgenic littermates were further obtained at ×10, × 63 magnification in the cortex and hippocampal areas and ×20 magnification for whole brain slices using a Leica SP8 confocal microscope (Leica Microsystems GmbH, Germany) at ScopeM. Sequential images were obtained by using 405 nm, 488 nm, and 561 nm lines. Identical resolution settings were used for the Z stacks (n = 15). FWHMs on the x- and y-axes were used for plaque size analysis using ×10 cortex confocal images for both the CRANAD-2 and 6E10 channels. The Allen brain atlas was used for anatomical reference (Jones et al., 2009).

## 3 Results

### 3.1 FMT reconstruction results and VOI analysis

A mesh containing 7065 nodes, 42021 tetrahedral elements and 1910 surface elements was generated for forward modeling, leading to a recovered fluorescence distribution stored in a Cartesian grid of 20 × 31 × 28. Overlays of FMT and MRI data are shown for the three perpendicular planes (**Fig. 2**). The structural information was clearly depicted by the T_2_*-weighted MR images, which helped localize the fluorescence signal significantly. The FMT signal displayed the progressive accumulation of the CRANAD-2 probe (by subtracting the baseline fluorescence intensity at t = 0 min) in the brain of the arcAβ mouse at t = 10, 20, and 30 min following the injection of the tracer (**Figs. 2a, b, c**). In all three cases, the CRANAD-2 probe had a higher concentration in the cortex region than in the subcortical region in a diffuse manner. VOI analysis in the cortical region showed that the uptake of CRANAD-2 increased continuously for 30 minutes. For the subcortical region (dashed line, **Fig. 2d**), the concentration of CRANAD-2 dropped after 20 minutes (**Fig. 2d**).

**Figure 2:**
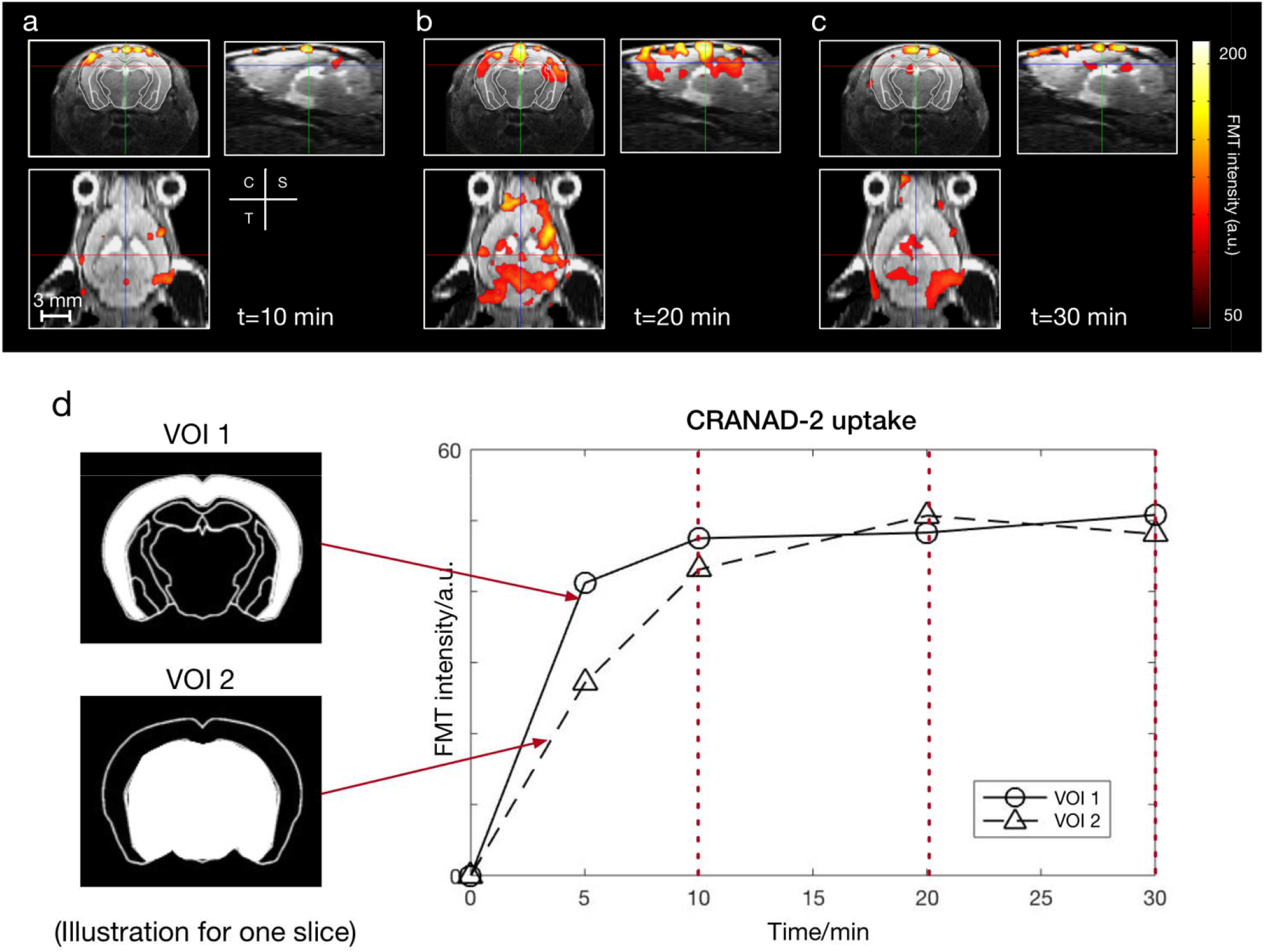
FMT reconstruction for visualizing CRANAD-2 distribution within the brain of one representative arcAβ mouse. FMT reconstruction shows variations at time points at t = 10, 20, and 30 minutes (a, b, c). VOIs at cortical and subcortical regions are indicated by two images of different masks in (d). The temporal variation in CRANAD-2 in the two VOIs is shown in (d). For the cortex (solid line), the uptake of CRANAD-2 increased for 30 minutes. For the subcortical region (dashed line), the concentration of CRANAD-2 dropped after 20 minutes.

### 3.2 Validation through FRI

FMT results were compared to data from two-dimensional FRI performed on another mouse under identical physiological conditions. The protocol of CRANAD-2 probe administration was identical to that used in the FMT experiment. Planar fluorescence images were recorded pre- and posti.v. injection of CRANAD-2 at the same 4 time points. Although no depth information was given with FRI, the fluorescent signal of CRANAD-2 showed the highest intensity at t = 20 minutes, which was similar to the result obtained with FMT (**Fig. 3**). However, FRI was not capable of disentangling the fluorescence distribution in the subcortex region due to the 2D nature of the imaging modality. The time course obtained from FRI showed a similar pattern as detected by using FMT.

**Figure 3:**
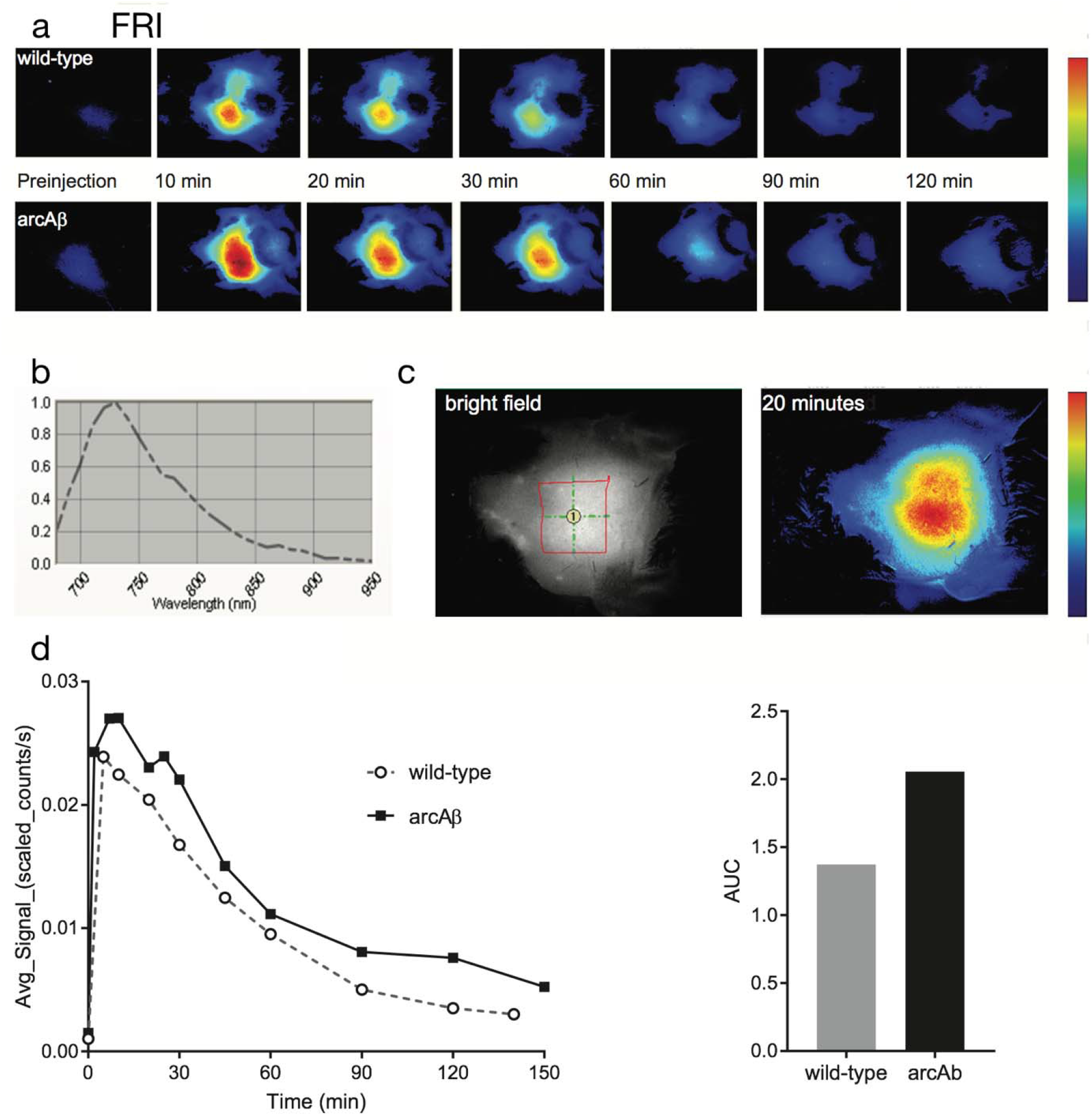
Time course analysis in fluorescence reflectance imaging (FRI) (a) Representative horizontal map of an NTL (upper row) and an arcAβ mouse (lower row) preinjection to 120 minutes after CRANAD-2 injection. (b) Spectra detected in the brains of arcAβ mice after injection; (c) White-light (left) and NIRF (right) images of one arcAβ mouse head with epilated skin over the head. Region of interest = 41 cm^2^; (d) Time course of NIRF signal in the brain of arcAβ and age-matched non-transgenic littermates; area under curve (AUC) of the cortical regions of arcAβ (n =4) and NTL (n =4) mice.

### 3.3 *Ex vivo* staining on mouse brain sections

To confirm the specificity of CRANDA-2 binding to Aβ deposits in the mouse brain, horizontal brain tissue sections from arcAβ mice and non-transgenic littermates were stained with CRANAD-2 in addition to thioflavin S and anti-Aβ antibodies 6E10 [65] and were nuclear counterstained with DAPI (Fig. 4). CRANAD-2 clearly co-stained with 6E10 and thioflavin S-stained parenchymal and vessel-associated Aβ deposits in the arcAβ mouse brain.

**Figure 4:**
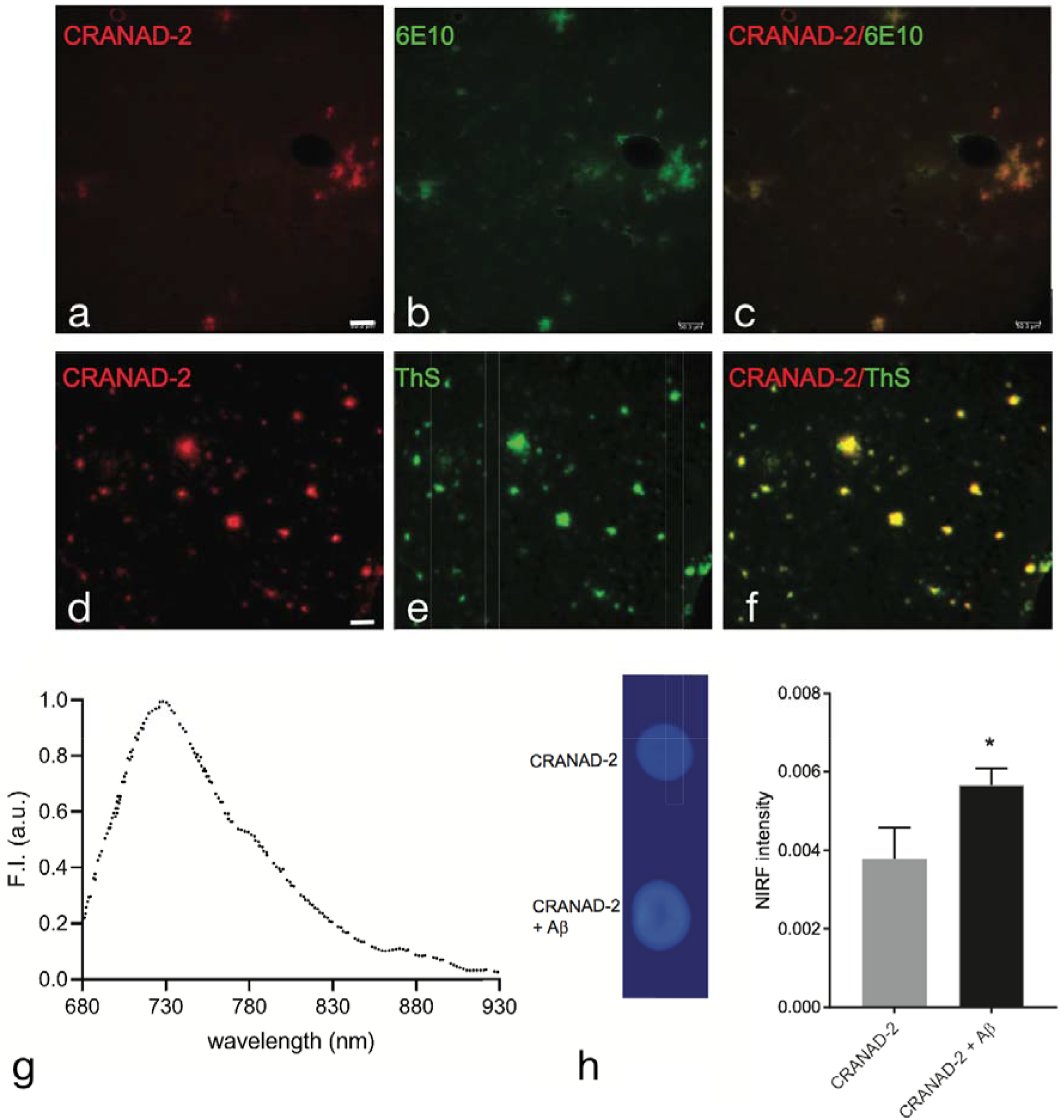
Co-staining of CRANAD-2 with Aβ deposition in non-transgenic littermate and arcAβ mouse brain tissue sections. (a-f) Sections were stained with CRANAD-2 (red) and anti-Aβ1-16 antibody 6E10 (green) in the cortex of arcAβ mice, and co-localization of the signal. (g) shows the emission spectrum of CRANAD-2 when binding with Aβ. Comparison of NIRF fluorescence signals with and without Aβ.

## 4 Discussion and conclusion

We have previously established a multimodal FMT-MRI framework assisted with the customized software platform STIFT that integrates the major steps during FMT reconstruction and optimizes its experimental configuration (Ren et al., 2020a). This framework can allocate the fluorescent signal to the registered high-resolution structural MRI reference regardless of whether the data have been acquired sequentially or simultaneously (Stuker et al., 2011). Furthermore, FMT reconstruction can incorporate a priori anatomic knowledge obtained by MRI to increase the accuracy of reconstruction. As the highly scattering nature of tissue renders FMT an ill-posed problem, implementing such prior information significantly improves the registration results (Arridge and Schotland, 2009). We have demonstrated the robustness and accuracy of the FMT-MRI method by using tissue-mimicking phantoms (Ren et al., 2020b) and in biological studies using a tumor-bearing mouse model (Ren et al., 2020a) or *ex vivo* intervertebral discs (Ren et al., 2021a).

In this work, local concentrations of the amyloid-fibril binding fluorescent probe CRANAD-2 were determined in the brains of aged arcAβ transgenic mice, which display a high load of cerebral amyloid plaques. The NIRF probe CRANAD-2 shows high-affinity binding and a redshift for Aβ aggregates and displays an increase in quantum yield upon binding with Aβ, thereby enhancing the signal-to-noise ratio. Sequential multimodal imaging combining FMT and MRI was applied to achieve VOI analysis guided by the MRI anatomical reference. After data acquisition, FMT-MRI image reconstruction and data analysis were conducted with the STIFT software platform (Ren et al., 2020a). A similar time course was observed from in vivo CRANAD-2 imaging using FMT and FRI, a routinely used planar imaging method. Strictly speaking, the mean fluorescence intensity value obtained from the region of interest (ROI) in FRI is based on 2D images, which is not equivalent to the 3D VOI values. Nevertheless, the FRI fluorescence signal close to the surface is highly weighted due to its reflection configuration. Thus, the comparison between the 2D ROI in the FRI and 3D VOI in the cortex region is reasonable.

There are substantial technical challenges associated with space constraints and compatibility of system components with the high magnetic field typically for preclinical MRI scanners. Nevertheless, the combination of FMT and MRI appears more attractive than FMT-CT for brain applications, given the high versatility of MRI and its superior soft tissue contrast compared to X-ray CT. Combined FMT-MRI studies have been reported for tumor imaging (Ren et al., 2016b). An early study allowed the differentiation of fibroadenoma in the breast of female patients from normal breast tissue based on differences in local perfusion/vascular permeability using concurrent contrast-enhanced MRI with a gadolinium (Gd) contrast agent and diffuse optical tomography with administration indocyanine green (ICG)(Ntziachristos et al., 2000). Similarly, dual-tracer MRI-guided fluorescence tomography was used to study the uptake kinetics of fluorescence dyes, which allowed estimation of the epidermal growth factor receptor (EGFR) density (Davis et al., 2013). Using a rapid image acquisition protocol that allowed dynamic monitoring of tracer uptake, tumor vascular perfusion and permeability was studied by simultaneous FMT-MRI measurements. In this case, fluorescent and MRI tracers were coinjected as a mixture, allowing head-to-head comparison of tracer uptake characteristics (Ren et al., 2016b). In all these aforementioned studies, postprocessing was rather time-consuming, as the whole procedure included registration of MR and optical images, sophisticated reconstruction using priors from MR structural images, and 3D visualization of the fluorophore distribution overlaid on the MRI images. In the present work, we successfully applied our customized software platform to efficiently tackle these steps. Our STIFT platform has the potential to further enhance the automation of FMT data acquisition by setting optimal parameters, such as the illumination pattern (Liu et al., 2019).

Imaging Aβ plaque distribution in the mouse brain using FMT is a challenging problem, as we have to consider complex anatomical structures such as the scalp and skull during reconstruction. The problem is further complicated by the fact that we do not consider focal brain lesions, e.g., in a glioma model (Ntziachristos et al., 2002), but rather a plaque distribution that is diffuse across cortical and, to some extent, subcortical structures. Thus, a low target-to-background ratio for the distribution of fluorescent dye across large brain areas is expected. Time-dependent binding of the Aβ-specific fluorescent probe CRANAD-2 was monitored in both cortical and subcortical regions over 30 minutes pre- and post-injection. The average intensity in the cortical region increased continuously, while the intensity in subcortical structures decreased at approximately t = 20 minutes after probe administration. This might be explained by the fact that amyloid-targeted CRANAD-2 accumulated in the cortex, whereas it was gradually washed out from regions not displaying the target, i.e., aggregated beta-amyloid proteins. The time point when the fluorescent signal reaches a maximum agrees well with the current and previous FRI using CRANAD-2 in an amyloidosis animal model (Ni et al., 2021; Ran et al., 2009).

However, we applied the multimodal FMT-MRI method in the AD study and achieved comparable results that agree well with the established FRI method. There are still some limitations to the current work. First, the data acquisition procedure for a single FMT measurement is now 5-10 minutes considering the point-by-point laser scanning illumination, which limits the temporal resolution of FMT. The optimization of illumination patterns and usage of structured light illumination can significantly reduce the scanning time and improve the temporal resolution of FMT (Liu et al., 2020; Ren et al., 2020a; Streeter et al., 2019). Second, in the current setup, FMT-MRI was implemented in sequential mode, i.e., the mouse was measured by standalone FMT and an MRI scanner independently. Taking the time for animal preparation and transport into account, we set the measurement time within 30 minutes. To more accurately study the kinetics of the CRANAD-2 probe, a longer measurement time (>30 min) will be preferred. This can be implemented by employing a hybrid FMT-MRI imager (Stuker et al., 2011), in which both MRI and optical images can be obtained simultaneously and the mouse can be measured for a longer time as no transport between different modalities is needed. Finally, FMT-MRI can also be compared with other emerging hybrid imaging methods, such as optoacoustic tomography-MRI hybrid systems, that can record concurrent optoacoustic and MR images with high spatiotemporal resolution (Ren et al., 2021b; Ren et al., 2019).

In conclusion, we demonstrated a non-invasive FMT-MRI multimodal strategy and a processing pipeline for *in vivo* imaging of cerebral Aβ deposits assisted with CRANAD-2 in AD amyloidosis mice. Such a pipeline may facilitate the understanding of the spread of amyloid deposits and the evaluation of Aβ clearing therapies.

## 6 Author Contributions

WR, RN, and MR conceived the study; WR and RN conceptualized the experiments. WR, RN, JK and MV performed the experiments; WR and JR programmed. WR and RN analysed the data. JK, MW and MR supervised the study. All authors contributed to writing and revising the manuscript.

## 8 Acknowledgements

The authors acknowledge support from Dr. Mark-Aurel Augath, Ms Gloria Shi, Dr. Marie Rouault at Institute for Biomedical Engineering, ETH & University of Zurich; Mr Daniel Schuppli at Institute for Regenerative Medicine, University of Zurich.

